# The ERBB network facilitates KRAS-driven lung tumorigenesis

**DOI:** 10.1101/290700

**Authors:** Björn Kruspig, Tiziana Monteverde, Sarah Neidler, Andreas Hock, Emma Kerr, Colin Nixon, William Clark, Ann Hedley, Craig Dick, Karen Vousden, Carla Martins, Daniel J. Murphy

## Abstract

KRAS is the most frequently mutated driver oncogene in human adenocarcinoma of the lung. There are presently no clinically proven strategies for treatment of KRAS-driven lung cancer. Activating mutations in KRAS are thought to confer independence from upstream signaling, however recent data suggest that this independence may not be absolute. Here we show that initiation and progression of KRAS-driven lung tumors requires input from ERBB family RTKs: Multiple ERBB RTKs are expressed and active from the earliest stages of KRAS driven lung tumor development, and treatment with a multi-ERBB inhibitor suppresses formation of KRas^G12D^-driven lung tumors. We present evidence that ERBB activity amplifies signaling through the core RAS pathway, supporting proliferation of KRAS mutant tumor cells in culture and progression to invasive disease *in vivo*. Importantly, brief pharmacological inhibition of the ERBB network significantly enhances the therapeutic benefit of MEK inhibition in an autochthonous tumor setting. Our data suggest that lung cancer patients with KRAS-driven disease may benefit from inclusion of multi-ERBB inhibitors in rationally designed treatment strategies.

**One Sentence Summary:** G12 Mutant KRAS requires tonic ERBB network activity for initiation and maintenance of lung cancer

## Introduction

Cancers of the lung account for over 1.5 million deaths per annum worldwide and 5-year survival rates remain between 10 & 15% in many developed countries [1]. The majority of lung cancers are classified as non-small cell (NSCLC) and adenocarcinoma is the most common histological subtype of NSCLC. Activating mutations in KRAS occur in a third of lung adenocarcinoma (LuAd) cases [2]. RAS proteins have historically proven to be elusive targets for selective inhibition, although the recent development of G12 mutant KRAS-selective tool compounds suggests that therapeutic targeting of KRAS may in time be possible [3, 4]. In the interim, there is a pressing need to develop alternative strategies for more effective treatment of KRAS-driven disease.

The ERBB family of receptor tyrosine kinases comprises 4 members, EGFR (ERBB1), HER2 (ERBB2, NEU), ERBB3 and ERBB4, all of which can homo- or heterodimerize, and dimerization is required for signaling activity. ERBB dimers are activated upon binding a spectrum of soluble ligands including EGF, Epiregulin, Amphiregulin and Neuregulin, amongst others, together forming a network for ERBB-driven signal transduction [5]. EGFR is a well-recognized driver of lung adenocarcinoma with genetic alterations present in up to 18% of cases [2]. ERBB2 and ERBB3 are highly expressed in embryonic lungs of humans and rodents and expression in both persists into adulthood [6, 7]. Overexpression of ERBB2 in the absence of gene amplification is common in human LuAd [8, 9] and functionality of ERBB2/ERBB3 heterodimers in NSCLC-derived cell lines was previously shown [10]. Amplification of any of the 4 ERBB RTKs is associated with poor prognosis in lung cancer [11], while high expression of the promiscuous ERBB ligand Epiregulin has previously been linked to disease progression and aggressive phenotypes in models of EGFR and KRAS-driven lung cancer [12, 13].

In a wild-type setting, ligand-activated signaling through ERBB RTKs activates KRAS [14]. Mutation of KRAS is generally thought to confer independence from upstream regulation, a view that is reinforced by the mutual exclusivity of activating mutations in KRAS and EGFR in LuAd, and by the failure of EGFR-selective inhibitors to show therapeutic benefit against KRAS-driven cancers [15, 16]. However, several recent results suggest that the independence of mutant KRAS from upstream signaling may not be absolute: In KRAS mutant NSCLC cell lines, activation of PI3K is contingent upon basal activity of wild-type IGFR, establishing an important precedent for coordination of oncogenic and normal signal transduction [17]; genetic deletion of EGFR was shown to suppress development of KRas^G12D^-driven pancreatic ductal adenocarcinoma [18, 19] while induced expression of ERBB2 and ERBB3 was found to underlie resistance of KRAS mutant lung and colorectal cell lines to MEK inhibition [20]. Strikingly, in the latter study, MEK inhibitor-induced ERBB2/3 expression was associated with recovery of phosphor-ERK levels downstream of KRAS, suggesting a surprising role for upstream signaling in sustaining pathway activity despite the presence of activated KRAS.

We therefore examined the requirement for ERBB activity in an inducible model of progressive autochthonous LuAd, driven by the combination of endogenously expressed KRas^G12D^ and modest overexpression of c-MYC. Here we present evidence that redundant signal transduction through multiple ERBB RTKs supports development and progression of mutant KRAS-driven lung tumors. Our data suggest that front-line use of multi-ERBB inhibitors may show clinical benefit in KRAS-driven LuAd.

## Results

### ERBB activity is required for KRas^G12D^-driven lung tumor formation

Induced expression of ERBB-family receptor tyrosine kinases (RTKs) is associated with resistance of KRAS mutant NSCLC cell lines to MEK inhibition [20]. We therefore examined expression of ERBB RTKs and their ligands in micro-dissected early-stage lung tumors, using a CRE-inducible mouse model of autochthonous lung adenocarcinoma driven by KRas^G12D^ combined with modestly elevated levels of MYC (lsl-KRas^G12D^;Rosa26-lsl-MYC – henceforth KM), the latter expressed from the Rosa26 locus at levels that alone fail to provoke a phenotype (Fig. S1)[21]. In tumor samples harvested 6 weeks after allele activation we found strong expression of Erbb2 and Erbb3 mRNA, while Egfr was weakly expressed and Erbb4 was not detected in tumors from 2 of 4 KM mice (Fig. 1A). Multiple ERBB ligands were expressed, with Areg, Tgfa and Hbegf showing strongest expression while Egf, Ereg, Nrg3 and Nrg4 were also clearly detected (Fig. 1B). The presence of both RTKs and multiple cognate ligands suggested that ERBB RTKs may actively signal in developing KRas^G12D^-driven lung tumors. Lysates prepared directly from multiple individual tumors, harvested 6 weeks post induction and immunoblotted for phosphor-ERBB RTKs, consistently showed readily detectable levels of phospho-Egfr, phospho-Erbb2 and phospho-Erbb3, suggesting that these RTKs are indeed active in developing KRas^G12D^-driven lung tumors (Fig. 1C). To determine if ERBB signaling functionally contributes to tumor development, we treated tumor-bearing mice acutely with a multi-ERBB inhibitor, Neratinib [22]. Treatment of mice for 3 days with Neratinib suppressed activity of ERBB RTKs, significantly reduced tumor cell proliferation and increased apoptosis, suggesting that ERBB activity promotes growth of KRas^G12D^-driven tumors (Fig. 1D-F). Strikingly, continuous daily treatment of mice from 2 weeks post allele induction almost completely suppressed the emergence of tumors, indicating that the combination of endogenously expressed KRas^G12D^ and constitutively expressed MYC requires input from ERBB RTKs in order to give rise to lung tumors (Fig. 1G, H). In contrast, daily Erlotinib treatment failed to show the same effect, consistent with reports that EGFR inhibition in isolation is ineffective in KRAS-driven lung cancer [15, 18].

**Figure 1:**
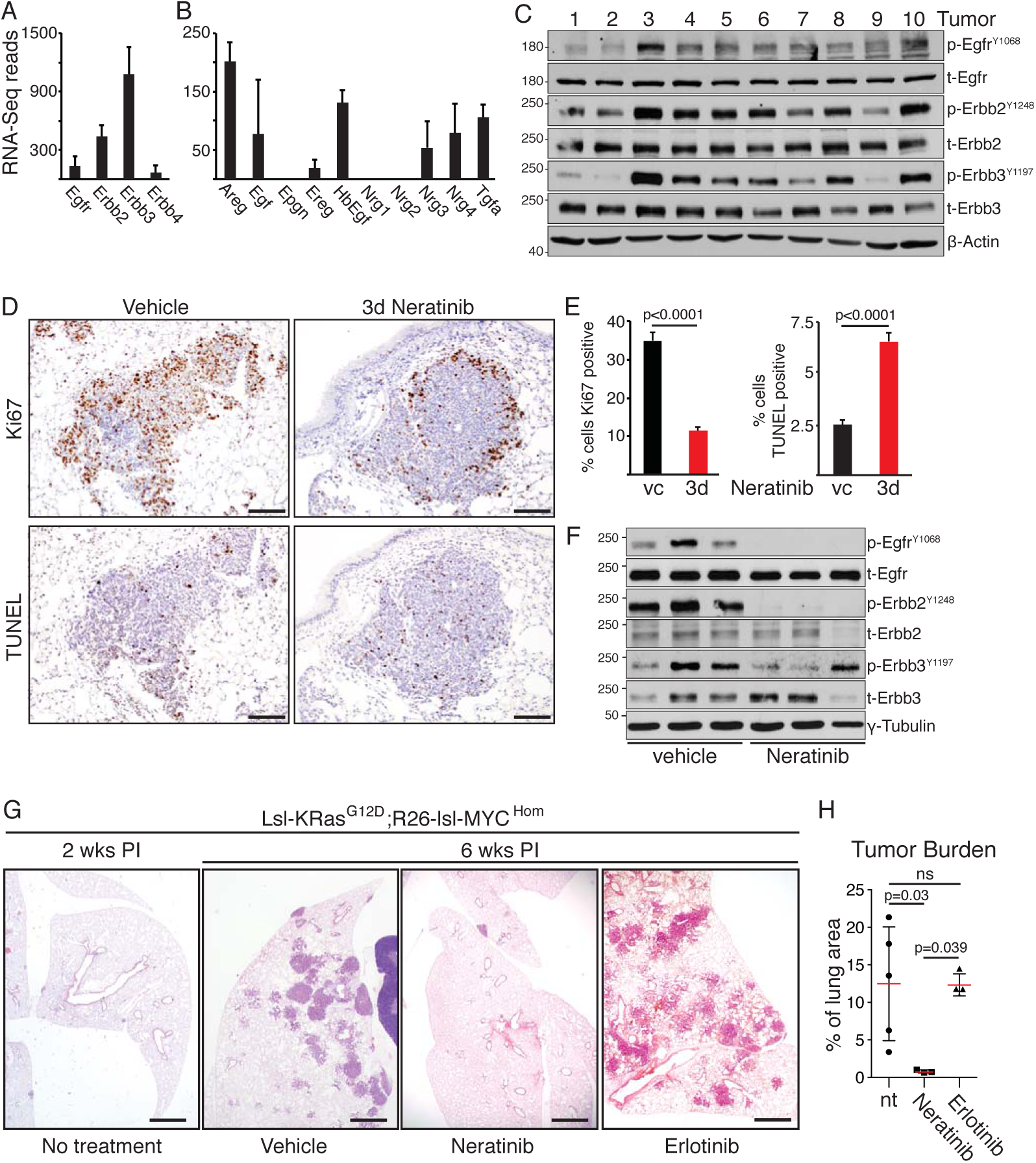
ERBB activity is required for KRAS-driven lung tumour formation. **A)** Expression of ERBB family RTKs in KM lung tumors harvested 6 weeks post allele induction (PI), measured by RNA-SEQ. Mean±SD read counts in tumours from 4 mice shown. **B)** Expression of ERBB family ligands in KM lung tumors harvested 6 weeks PI, as per (A). **C)** Immunoblots of lysates generated from 10 individual KM tumors, harvested 6 weeks PI, using the indicated antibodies. **D)** Representative images of IHC for Ki67 and TUNEL staining of KM mice treated for 3 days with Neratinib (N=3) or vehicle control (N=3). Scale bars = 100µm. **E)** Quantification (Mean ± SEM) of staining in 5 tumours from each mouse as per (D). Left panel shows % of tumor cells expressing Ki67; right panel shows % of TUNEL positive tumor cells; vc = vehicle control. P values are from 2-tailed T tests. **F)** Immunoblots of 3 individual KM tumors from mice treated for 3 days with Neratinib or vehicle control. **G)** Representative H&E images from KM mice treated daily as indicated, commencing 2 wks PI, and harvested at 6 wks PI. Scale bar = 1mm. **H)** Quantification of tumor burden from (G). Each data point represents the mean value across 3 sections, separated by 100µm, from each mouse. ANOVA followed by Tukey test. Ns=not significant.

### Progression of KRas-driven lung tumors is associated with increased ERBB network expression

The above analysis was performed on tumors at 5-6 weeks post Adeno-CRE-mediated allele activation. At this time post induction, the lungs of all induced KM mice contain dozens of individual tumors that present with uniform histology resembling human papillary adenocarcinoma in situ (Fig. 2A, panel i). A small minority of tumors (2-5%) also contain a second, more disorganized population with more aggressive histological features, including increased cytosolic and nuclear volume, prominent nucleoli, and increased morphological heterogeneity (Fig. 2A, panel ii). IHC analysis revealed continuous expression of the same lineage markers across both populations in individual tumors, suggesting that the second populations represent the emergence of aggressive sub-clones (Fig. S1B). Progression of KRas^G12D^-driven lung tumors to more aggressive disease is associated with a pronounced increase in Erk1/2 phosphorylation [23, 24]. Accordingly, IHC analysis of phospho-Erk (p-Erk) levels revealed sharply higher expression of p-Erk in these disorganized sub-clones compared with the rest of the same tumors, and indeed across the entire tumor population (Fig. 2A & B). By comparison, KM tumors harvested at 4-6 months post-induction showed widespread expression of p-Erk, as these more aggressive sub-clones gradually come to predominate (Fig. 2B, panel iii). Moreover, KM metastases to the liver also show high levels of epithelial p-Erk staining (Fig. 2C).

**Figure 2:**
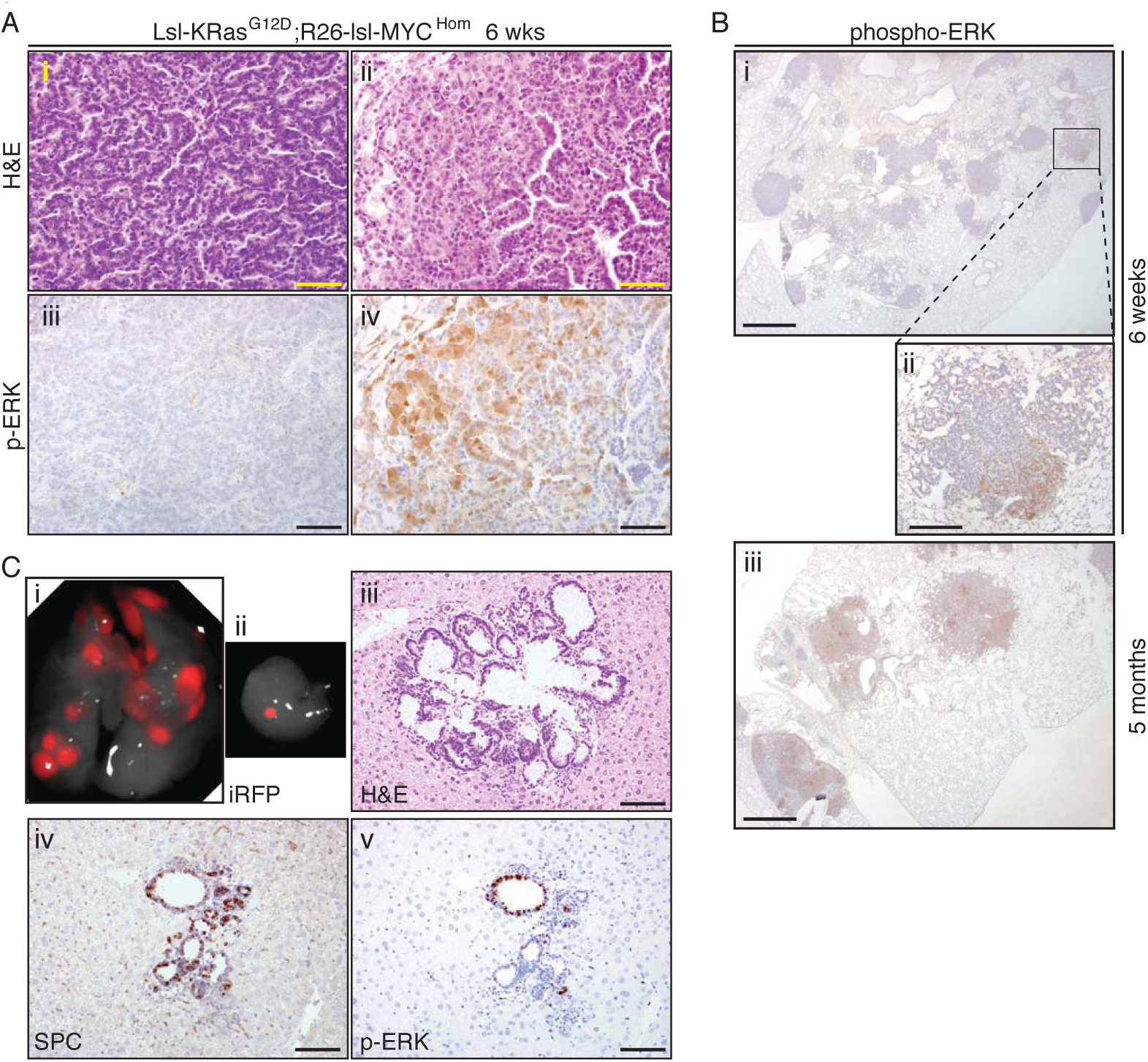
KM lung tumor progression is associated with increased ERK phosphorylation. **A)** Images of H&E (upper panels) and phospho-Erk (lower panels) stained KM lung tumors harvested at 6 wks PI illustrating histological changes associated with tumor progression: Panels i & iii are representative of >95% of KM lung tumor area at 6wks; panels ii & iv are representative of 2-5% of total tumor area at 6 wks. Scale bar = 50µm. **B)** Phospho-ERK staining in KM tumors harvested at 6wks PI (i & ii) versus 5 months PI (iii). Scale bars = 1mm (i & iii) and 200µm (ii). **C)** Primary lung tumors (i) and dissected liver fragment with metastasis (ii-v) in a KM mouse harvested 6 months PI. Panels i & ii show fluorescent detection of a *Hprt-lsl-IRFP* CRE-reporter allele induced concomitantly with *R26-lsl-MYC^DM^* and *lsl-KRas^G12D^* alleles by intra-nasal Adeno-CRE installation (same magnification). Also shown: H&E (iii), SPC (iv) and p-Erk (v) staining of liver metastasis. Scale bar = 50µm. Images are representative of 3 mice.

Increased p-Erk levels were previously shown to be associated with amplification of the mutant KRas locus in a KP lung cancer model wherein KRas^G12D^ expression was combined with loss of functional p53 [25]. We used laser-capture micro-dissection coupled with RNA-SEQ analysis to compare gene expression in p-Erk^Low^ with p-Erk^High^ KM tumor regions from 4 individual mice (see schematic, Fig. S1C). Expression of KRas was modestly higher (< 2 fold) in pErk^High^ tumor regions, suggesting that locus amplification is not the underlying driver of progression in the KM model (Fig. 3A). Strikingly, we detected a sharp increase in expression of promiscuous ERBB-family ligands, Epiregulin (Ereg) and Amphiregulin (Areg), along with more modest but significant increases in HbEgf and Tgfa (Fig. 3B-D and Fig. S1D). Elevated expression of Ereg and Areg were previously reported to be associated with more aggressive tumors in a titrated model of KRas-driven mammary cancer [26] and in human NSCLC cells enriched for metastatic behavior [27]. In situ hybridization revealed a clear overlap between expression of Ereg, Areg and high levels of p-Erk (by IHC), confirming the RNA-SEQ data (Fig. 3C). Additionally, RNA-SEQ analysis revealed significantly increased expression of Adam9 & Adam10 sheddases, which process membrane-bound ERBB ligands into their more potent soluble forms [28]; increased expression of the signaling scaffold Iqgap2; and increased expression of LamC2, Fabp5 and Keratin19, all recently shown to enhance signaling through EGFR/ERBB family RTKs [27, 29, 30]. Expression of Erbb2 and Erbb3 also increased in p-Erk^High^ regions, albeit not quite reaching statistical significance (Fig. 3D & E).

**Figure 3:**
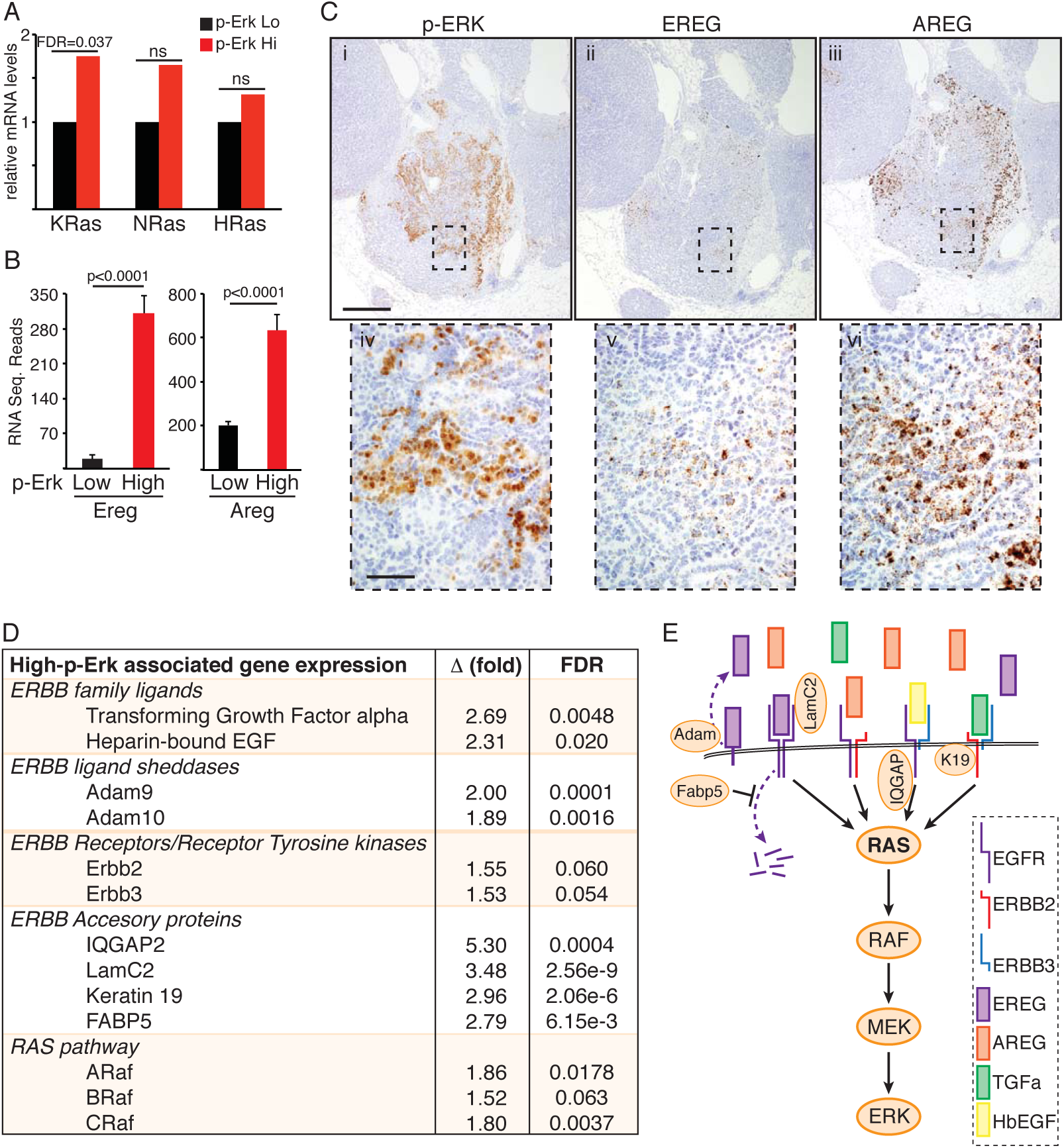
Increased expression of the ERBB network during progression from p-Erk^low^ to p-Erk^high^ KM tumors. **A)** Normalised expression of RAS genes in laser-capture micro-dissected (LCM) p-Erk^High^ KM tumor regions relative to p-Erk^Low^ regions from tumors in the same mice (N = 4 mice), measured by RNA-SEQ. False discovery rate (FDR) shown for KRAS; ns = not statistically significant. **B)** Mean and SEM RNA-SEQ reads of Ereg and Areg mRNA from p-Erk^Low^ & p-Erk^High^ KM tumor regions from 4 mice. Adjusted p values were calculated in R. **C)** Serial sections of KM tumors stained by IHC for p-Erk (i, iv) or by in situ hybridization for Ereg (ii, v) or Areg (iii, vi). Scale bars = 200µm (panels i-iii) & 25µm (panles iv-vi). **D)** Normalised expression of ERBB network genes showing mean fold increase in expression in p-Erk^High^ relative to p-Erk^Low^ KM tumor regions from 4 mice as per (A). FDR = false discovery rate. **E)** Diagramatic representation of significantly up-regulated components of the ERBB-RAS-ERK pathway associated with KM tumor progression.

These data suggest an alternative route to increased RAS pathway signaling that does not require KRas amplification but rather involves increased activity of the ERBB network, defined here as comprising ERBB ligands, RTKs and RTK accessory molecules (eg. LamC2 & Fabp5). Notably, examination of the TCGA lung adenocarcinoma (LuAd) dataset via cBioPortal [31] revealed overexpression and/or amplification of one or more constituents of the ERBB network in the majority of KRAS mutant human LuAd (Fig. S2). By comparison, amplification of the mutant KRAS locus occurs in a rather smaller proportion of KRAS mutant LuAd and amplification shows a tendency towards mutual exclusivity with ERBB ligand overexpression.

### ERBB signaling amplifies RAS pathway activity in KRAS mutant human NSCLC cells

Combined inhibition of both EGFR and ERBB2 is required to prevent outgrowth of MEK inhibitor-resistant clones of human KRAS mutant NSCLC cells, however the effect of multi-ERBB inhibition on treatment-naïve cells was not previously explored [20]. Treatment of multiple KRAS mutant human NSCLC lines with the multi-ERBB inhibitor Neratinib suppressed proliferation in a dose-dependent manner (Fig. 4A). In contrast, inhibition of EGFR in isolation showed little effect. Immunoblotting confirmed that the doses of Neratinib used strongly suppressed activity of ERBB RTKs (Fig. 4B). Consistent with previous reports that Neratinib drives increased ERBB turnover [32], ERBB2 and ERBB3 protein expression were also reduced by Neratinib in multiple cell lines. Despite the presence of G12 mutant KRAS in all such cells, Neratinib consistently reduced levels of phosphor-ERK, suggesting that ERBB signaling amplifies RAS pathway activity even in cells expressing mutant KRAS (Fig. 4B & S3A). We therefore examined RAF binding as a direct measure of RAS signaling activity in A549 cells that are homozygous for mutant KRAS. Acute ERBB inhibition reduced RAS:RAF binding by some 50%, consistent with the observed partial reduction in p-ERK levels (Fig. 4C). Additionally, murine lung tumor cells carrying a spontaneously amplified KRas^G12D^ allele also showed sensitivity to Neratinib, albeit somewhat lower than that of cells carrying a single copy of the allele (Fig. S3B). Together these results suggest that sensitivity to ERBB suppression is imparted through both mutant and wild-type KRAS.

**Figure 4:**
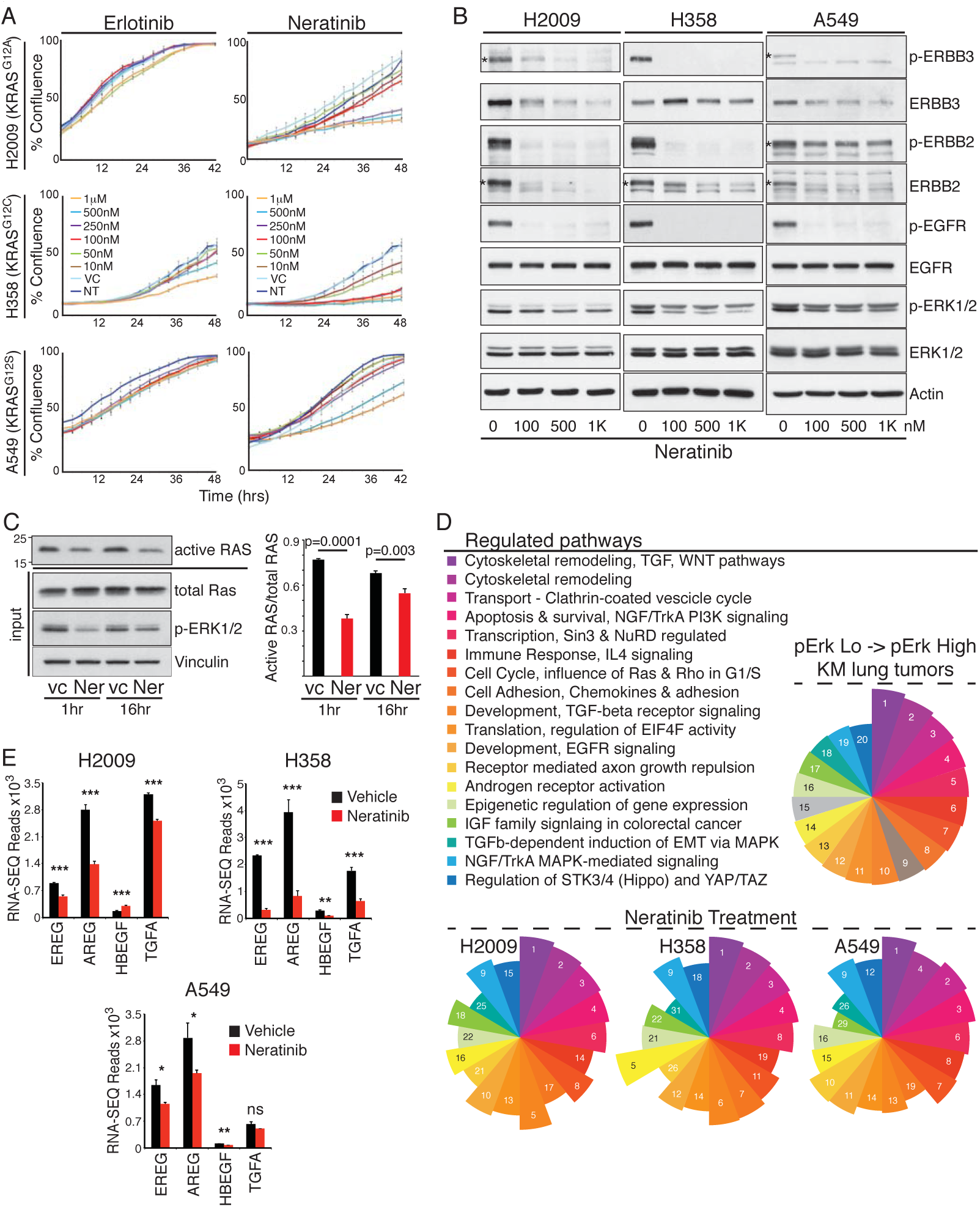
A feed-forward ERBB signalling loop drives proliferation of KRAS mutant human NSCLC cells. **A)** Growth curves of 3 KRAS mutant human NSCLC lines upon treatment with increasing doses of the EGFR-selective inhibitor Erlotinib or the dual EGFR/ERBB2 inhibitor, Neratinib, measured by Incucyte time-lapse video-microscopy. Error bars show SD for technical triplicates. Data are representative of at least 2 independent experiments. **B)** Lysates from KRAS mutant NSCLC cells treated with increasing doses of Neratinib, immunoblotted with the indicated antibodies. Asterisks, where present, indicate the correct band. **C)** RAS immunoblot of RAF-coated bead precipitates (top panel) from lysates of A549 cells treated with Neratinib or vehicle control for 1 or 16 hrs. Lysate input aliquots were immunoblotted with the indicated antibodies. Right panel shows Mean±SEM quantification of band intensities from 3 independent experiments (arbitrary units). P values are from 2-tailed T-tests. **D)** Top 20 significantly modulated pathways associated with the transition to p-Erk^High^ disease in the KM model, identified using Metacore GeneGO analysis of RNA-SEQ expression data. Segment size in the pie-chart reflects ranking of the pathways by false discovery rate (FDR). Lower panel shows 18 of the same pathways are significantly modulated in each of 3 KRAS mutant human NSCLC lines after overnight treatment with Neratinib. Numbers and pie segment size reflect ranking by FDR. **E)** Expression of ERBB ligands in the indicated cells treated overnight with vehicle (black) or Neratinib (red), measured by RNA-SEQ as per (D). Mean & SEM of biological triplicates shown. P values are from 2-tailed T-tests, abbreviated as follows: * =<0.05; ** = <0.01; *** <0.0001; ns = not significant.

To gain further mechanistic insight into the growth inhibitory effects of Neratinib, we performed unbiased transcriptomic analysis after overnight treatment of 3 KRAS mutant NSCLC cell lines. Metacore GeneGO pathway analysis revealed a remarkable degree of consistency across the 3 cell lines tested. Most strikingly, 18 of the 20 most highly modulated pathways associated with progression of murine KM tumors to p-Erk^High^ disease were reciprocally modulated in the human cells upon Neratinib treatment (Fig. 4D & Table S1). Significantly, in all three cell lines, ERBB inhibition reduced expression of the same ERBB ligands that increased as KM tumors progress to p-Erk^High^ disease. Taken together, ERBB activity thus establishes a feed-forward loop that sustains KRAS mutant NSCLC proliferation *in vitro* and drives tumor progression *in vivo*, at least in part by amplifying signaling through the core RAS-ERK module.

### ERBB inhibition enhances the potency of MEK inhibition *in vitro* and extends survival of mice with LuAd

With the exception of H358 cells, ERBB inhibition did not alone result in death of KRAS mutant NSCLC cells *in vitro*, however, Neratinib significantly increased apoptosis induced by inhibition of MEK downstream of KRAS in multiple cell lines (Fig. 5A). Substitution of Neratinib with a second multi-ERBB inhibitor, Afatinib, perfectly replicated the effects of Neratinib, confirming the on-target specificity of the drug (Fig. S3C, D). Moreover, both drugs combined with MEK inhibition to suppress colony formation (Fig. 5B). Consistent with published results [20] MEK inhibition alone increased expression of ERBB2 and ERBB3 *in vitro*, although the effect on ERBB2 phosphorylation was variable across the cell lines tested. Importantly, co-treatment with Neratinib continued to suppress both expression and activity of ERBB2 and ERBB3, while also suppressing EGFR activity (Fig. 5C).

**Figure 5:**
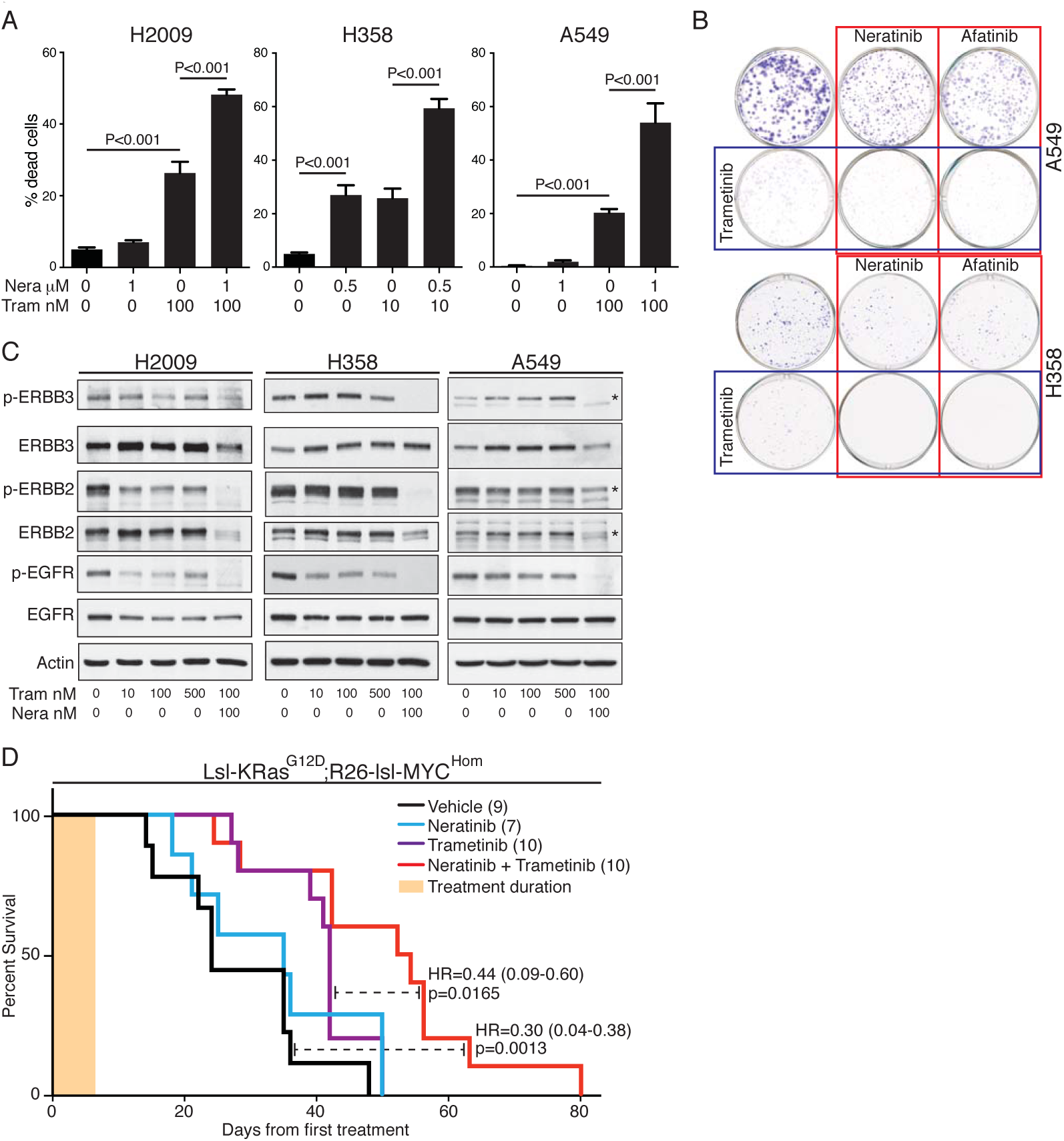
ERBB blockade enhances MEK inhibitor-driven apoptosis *in vitro* and therapeutic impact *in vivo*. **A)** Apoptosis induced in human NSCLC cells, measured 48hrs after treatment with the indicated doses of Neratinib (Nera) and/or Trametinib (Tram). Mean ± SEM of 3 independent experiments shown. ANOVA followed by Tukey test. **B)** Clonogenic assay showing suppression of colony formation in A549 and H358 cells after 48hr treatment with the indicated inhibitors. **C)** Lysates from the indicated cells treated for 24hrs with increasing doses of Trametinib alone or the combination of Trametinib and Neratinib, immunoblotted with the indicated antibodies. Asterisks, where present, indicate the correct band. **D)** Overall survival, measured from the first day of treatment, of tumor-bearing KM mice treated daily for 1 week (tan bar) with Neratinib (80mg/kg), Trametinib (1mg/kg) or the combination of both, then followed without further intervention. Treatment was commenced at 5 weeks PI. Cohorts shown are vehicle (N=9); Neratinib (N=7); Trametinib (N=10); Trametinib + Neratinib (N=10). Logrank hazard ratios (HR±95% CI) & p values are shown for comparisons of T+N versus vehicle and T+N versus T alone (dashed lines).

The combination of MEK and multi-ERBB inhibition was previously shown to suppress the growth of sub-cutaneous NSCLC xenografts under continuous daily treatment [20], however, cell lines xenografts have a poor track record of accurately predicting human patient responses. We therefore tested the therapeutic efficacy of transient inhibition of ERBB and/or MEK in our fully immuno-competent KM model of autochthonous lung adenocarcinoma. Tumors were induced in adult KM mice and allowed to develop for 6 weeks. The presence of an IRFP Cre-reporter allele [33] in a subset of such animals allowed us to confirm the presence of tumors by Licor PEARL fluorescence imaging prior to initiating therapy (Fig. S4). Mice were treated daily for 1 week with Neratinib, alone or in combination with Trametinib, then left untreated to determine the effect on overall survival. Transient ERBB blockade alone showed little influence on overall survival whereas MEK inhibition alone significantly extended survival of KM mice. Importantly, the combination of Neratinib and Trametinib further extended survival significantly over that achieved by MEK inhibition alone (Fig. 5D). Fluorescence imaging of IRFP positive KM mice showed pronounced suppression of lung tumor growth in individual mice treated with combination therapy (Fig. S4). We conclude from these data that multi-ERBB inhibition may benefit LuAd patients with mutant KRAS-driven disease, if used in combination with other agents such as MEK inhibitors.

## Discussion

EGFR-selective inhibitors have failed to show clinical benefit in mutant KRAS-driven cancers. In contrast with targeted inhibition of EGFR in isolation, we show here that broad inhibition of the ERBB network, using the multi-ERBB inhibitor Neratinib, almost completely suppresses formation of KRas^G12D^-driven lung tumors. We show that ERBB activity enhances signaling though the core RAS->ERK pathway, establishing a feed-forward loop that is associated with progression to invasion and metastasis. Importantly, we show that brief, transient inhibition of ERBB signaling significantly enhances the therapeutic benefit of MEK inhibition in the autochthonous setting. Although this latter result is broadly supportive of the clinical implications of a prior study showing that increased ERBB activity underlies resistance to MEK inhibition [20], our data are fundamentally distinct in that we demonstrate a clear role for ERBB activity *from the very outset* of KRAS-driven tumor initiation, as opposed to in reaction to targeted inhibition of MEK.

In the course of preparing this manuscript we became aware of an independent study that strongly complements our observations and underscores our conclusions: The work by Moll and colleagues similarly demonstrates a requirement for ERBB signaling to support progression of KRas^G12D^-driven lung cancer *in vivo* [34]. Importantly, their study utilized an independent pan-ERBB inhibitor, Afatinib, in the context of both KRas^G12D^-only and KRas^G12D^;p53^Fl/Fl^-driven tumor models. The striking similarities in the two studies attests to the on-target specificity of the 2 drugs used while also showing that the effects are independent of the genetic strategy used to accelerate KRas-driven disease and are therefore likely to have broader potential application.

Previous work has revealed that threshold levels of KRAS signaling are required for tumor initiation and progression. In a titratable model of HRas^G12V^ overexpression in mammary glands, focal tumors arising spontaneously from non-transformed epithelium that expressed the transgene at low levels exhibited a pronounced increase in expression of both the HRas^G12V^ transgene and of multiple ERBB ligands, including Ereg, Tgfa and HbEgf, again suggestive of feed-forward signal amplification [26]. More recently, this same “activity threshold” principle was shown to apply to MAPK (ERK) signaling as the key oncogenic effector pathway of KRAS in the lung [35]. Through pharmacological enhancement of MAPK activity, this study showed that different airway cell types require distinct levels of MAPK signaling for oncogenic transformation by mutant KRas, and that a second, higher, threshold signal was needed for progression to carcinoma. The existence of this higher threshold in KRas-driven LuAd was already clear from the pronounced increase in phosphor-Erk levels associated with tumor progression [26] which, in the KP model, is driven by spontaneous amplification of the G12D mutated allele [25]. Our data suggest that ratcheting up ERBB signal transduction provides an alterative route to RAS *pathway* signal amplification, independent of *KRAS* gene amplification. This is supported by our analysis of human data that shows a trend towards mutual exclusivity of ERBB ligand overexpression and amplification of mutant KRAS. It is important to note that, from the human data, there does not appear to be a preferred mechanism of ERBB signal enhancement – overexpression of ligands, amplification of RTKs and/or accessory molecules could all plausibly achieve the same effect. What is clear from these data is that the potential to increase ERBB signaling appears to be widespread in KRAS mutant human LuAd.

Perhaps the most surprising observation is that endogenously expressed KRas^G12D^ requires basal signaling from ERBB RTKs in order to initiate lung tumors, even when accompanied by MYC overexpression. Note that we previously showed that expression from the Rosa26 locus is refractory to growth factor signaling [36] – thus the requirement for ERBB activity does not reflect an artificial need for ERBB to sustain expression of the MYC transgene. Instead, these data suggest that G12 mutant KRAS requires a push from upstream RTKs in order to breach the initial threshold needed for tumor initiation. Supporting this hypothesis, it was recently shown that G12 mutant KRAS, although no longer responsive to RAS-GAPs, does retain some level of intrinsic GTPase activity and thus cycles slowly between on and off states [4], opening the possibility for upstream signaling to influence activity, either directly or by promoting activity of RAS guanonucleotide exchange factors (GEFs). Alternatively, tonic signal transduction through wild-type RAS isoforms may need to combine with that from mutant KRAS to likewise breach the threshold for transformation [37]. Our data presently do not distinguish between these possibilities and, indeed, they may not be mutually exclusive.

Activating mutations in KRAS are currently considered grounds for exclusion from clinical treatment with EGFR/ERBB inhibitors, while single-agent trials of the same drugs have failed to show efficacy outwith of cohorts with activating mutations in EGFR or ERBB amplification [38]. Indeed, our data concur that multi-ERBB inhibitors are unlikely to benefit KRAS-driven cancer patients if used in isolation. However, we show here the potential for such drugs to sensitize autochthonous KRAS-driven tumors to additional therapeutic agents, in this case to the MEK inhibitor Trametinib. Our data argue that clinical use of multi-ERBB inhibitors as part of an inhibitor cocktail to treat KRAS-driven LuAd deserves re-examination.

## Materials and Methods

### Genetically Engineered Mice & Mouse Procedures

Procedures involving mice were performed in accordance with Home Office license numbers 60/4183 & 70/7950 (CRUK BICR, UK). *LSL-KRas^G12D^* mice [39] were obtained from the NCI mouse repository at Fredrick, MD, USA. Rosa26^DM.lsl-MYC^ mice were generated as previously described [36] with the ATG-initiated human *c-MYC* (MYC2) cDNA, devoid of 5’ and 3’ UTRs, replacing the MYC-Estrogen Receptor ligand-binding domain fusion cDNA. Targeted insertion into the *Rosa26* locus was confirmed by Southern blotting and genotyping was initially performed using the following primers: A) CCC AAA GTC GCT CTG AGT TG (common); B) GCG AAG AGT TTG TCC TCA ACC (targeted locus); C) GGA GCG GGA GAA ATG GAT ATG A (wild-type locus). All genotyping was subsequently performed by Transnetyx Inc.. A more detailed description of this mouse is the subject of a separate manuscript, in preparation at the time of writing [21]. The Hprt-lsl-IRFP allele was previously described [33]. Mice were maintained on a mixed FVBN/C57Bl6 background, housed on a 12hr light/dark cycle and fed and watered ad libitum. Recombinant Adenovirus expressing Cre was purchased from the University of Iowa gene therapy facility. For Adeno-Cre installation, young adult (8-10 week old) mice were sedated with a mixture of Domitor and Ketamine, injected IP. For most experiments, 1×10^7^ pfu Adeno-CRE were administered intranasally using the calcium phosphate precipitation method, as described previously [36]. For lower tumor burden, 3×10^5^ pfu were administered. For survival analysis, impaired breathing and/or hunching with no more than 10% weight loss was chosen as an endpoint. Neratinib (Medchem Express) was administered by twice-daily gavage of 40mg/kg for up to 4 weeks, or a single daily IP injection of 80mg/kg for short-term (3-7 days) experiments. For gavage, 0.5% methylcellulose, 0.4% Tween-80, in H_2_O was used as vehicle; peanut oil was used for IP. Trametinib was administered in peanut oil at 1mg/kg. Live imaging of iRFP positive tumor-bearing mice was performed using a PEARL imaging system (Licor). All mice were sacrificed humanely using a schedule 1 procedure.

### Immunohistochemistry and Tissue Analysis

Mouse tissues were perfusion fixed in zinc formalin overnight. 4µm paraffin sections were de-paraffinized and rehydrated: 3 × 5 minutes xylene, 2 minutes in each 2×100%; 2×95%; 2×70%; 1×50% ethanol; dH_2_O. Peroxidase blocking was performed for 10 min in 3% H_2_O_2_ diluted in H_2_O, followed by antigen retrieval in 10mM citrate buffer, pH 6, 10 min near boiling by microwave heating at low power. Non-specific antibody binding was blocked with up to 3% BSA or up to 5% normal goat serum for 1h at RT or overnight at 4°C. Primary antibody incubation was performed overnight at 4°C or 2h at 37°C. Secondary biotinylated antibody was incubated 1h at room temperature and stain was developed with stable DAB (Invitrogen) followed by counterstaining with Gil 1 hematoxylin (Sigma MH216) and Scotts tab water substitute. The following antibodies were used at the indicated dilution: p-Erk (P44/42 MAPK phosphor-Thr202/Tyr204), Cell Signaling CS4370, 1:500; Ki67 (Sp6), Fisher Scientific RM9106S 1:200; SP-C, Millipore AB3786, 1:1000; CC10, Millipore 07-623, 1:1000; TUNEL ApopTag kit, Millipore S7100; Vectorlabs VECTASTAIN ABC kit; anti-Rabbit IgG, PK-4001 ECL; anti Rat, GE Healthcare NA935. Quantification of Ki67 and TUNEL labeling was scored manually on 5 tumors from each mouse and graphs show mean values ± SE from the indicated number of mice. Tumor burden was determined using Leica software as the % area of lung tissue occupied by tumors, measured on 3 Hematoxylin & Eosin (H&E) stained sections, separated by at least 100mm, from each of the indicated numbers of mice. Histological classification of tumours as adenocarcinoma was determined independently by 2 clinical pathologists.

### Laser-Capture Microdissection & RNA-SEQ analysis

For micro-dissection, 10mm FFPE sections were mounted on framed membrane slides (Leica). Adjacent sections were mounted on standard poly-L-lysine-coated glass slides and stained for p-Erk. Membrane slide-mounted tissue was de-paraffinized, rehydrated and stained with ice-cold 1% cresyl violet. Selected tissue was micro-dissected using a Leica DM 6000B microscope and total time for staining and micro-dissection was under 20 minutes per sample. P-Erk^Low^ and p-Erk^High^ tumor regions were harvested into separate tubes and tissue from multiple sections was pooled for each of 4 mice. Harvested tissue was suspended in 30ml PKD buffer (Qiagen, RNEasy FFPE kit) and flash frozen for storage. Total tissue RNA was isolated using the RNEasy FFPE kit according to manufacturer’s directions and ribosomal RNA was depleted with Ribo-Zero (Epicentre). Synthesis of cDNA was performed using the SMARTER Stranded random primed RNA-SEQ kit (Takara/Clontech), resulting in cDNA libraries flanked by Illumina indexing primers. Following library quantification (Quant-IT Pico green kit, Invitrogen), libraries were standardized to 10nM, denatured, diluted to 10pM and analyzed by paired-end sequencing using an Illumina GA11X deep sequencer. The raw RNA-sequencing data files underwent quality checks using FastQC and FastQ-Screen software. RNA-sequencing reads were aligned to the GRCm38 [40] version of the mouse genome using Tophat2 version 2.0.10 [41] with Bowtie version 2.1.0 [42]. Relative expression levels were determined and statistically analyzed by a combination of HTSeq and the R 3.0.2 environment, utilizing packages from the Bioconductor data analysis suite and differential gene expression analysis based on the negative binomial distribution using the EdgeR package [43]. Pathway modulation analysis was performed using Metacore GeneGO (Thompson Reuters). For analysis of individual gene expression, data were normalized internally to B2M for each mouse, and fold change and false discovery rates were recalculated. RNA-SEQ analysis of human cell lines: Total RNA was isolated using the RNEasy Mini Kit (Qiagen) according to manufacturer’s instructions and DNA was depleted with the RNase-Free DNase Set (Qiagen). RNA-integrity was checked using the RNA ScreenTape assay (Agilent Technologies) and cDNA was synthesized with the TruSeq Stranded mRNA Library Prep Kit (Illumina). Following library quantification (D1000 ScreenTape, Agilent Technologies), libraries were standardized to 10nM, denatured, diluted to 10pM and analyzed by paired-end sequencing using an Illumina NextSeq500 platform. RNA-Sequencing reads were aligned to the GRCh37 version of the human genome and differential expression determined using DESeq2 [44].

### Cell Culture and Related Assays

Human lung cancer cell lines (A549, H2009, H358) were validated in-house and grown in RPMI with 10% FBS & penn/strep. For immunoblotting, cells were lysed in RIPA^Hi^ buffer (150mM NaCl, 50mM Tris [pH 7.5], 1% NP-40, 0.5% sodium deoxycholic acid, 1% SDS plus cOmplete protease/phosphatase inhibitor cocktails [Sigma-Aldrich]) and after western blotting probed with the following primary antibodies: p-ERK Thr202;Tyr204 (E-4), Santa Cruz SC7383; p-EGFR Tyr1068, Cell Signaling 3777; p-ERBB2 Tyr 1248, Millipore 06-229; p-ERBB3 Tyr 1197 Cell Signaling 4561; ERK1/2 Cell Signaling 4695; EGFR Millipore 06-847; ERBB2 Merck OP15L; ERBB3 Millipore 05-390. RAS:RAF binding assays were performed using a commercial RAS activity assay kit (Cytoskeleton). Cell propagation and death was analyzed by Incucyte time-lapse video-microscopy in the presence of Sytox green. Cell death measurements were performed after 48hrs drug treatment and corrected for confluence. All experiments were performed in biological triplicate, except where noted.

### Statistical Analysis

Raw data obtained from quantitative Real Time PCR, FACS and Incucyte assays were copied into Excel (Microsoft) or Prism (Graphpad) spreadsheets. All Mean values, SD, and SEM values of biological replicates were calculated using the calculator function. Graphical representation of such data was also produced in Excel or in Prism. Statistical significance for pairwise data was determined by the Student’s T test. For multiple comparisons, ANOVA was used with a post-hoc Tukey test. * denotes P<0.05; ** denotes P<0.01; *** denotes P<0.001. For Kaplan-Meier plots Mantel Cox logrank P values are presented.

## Supplementary Materials

Fig. S1. Characterization of KM lung tumors. Relates to main figures 1, 2 & 3.

Fig. S2. Genomic alterations and expression data of ERBB network genes in human KRAS mutant Lund Adenocarcinoma. Relates to main figure 3.

Figure S3. Figure S3: ERBB blockade enhances MEK inhibitor induced NSCLC cell death. Relates to main figures 4 & 5.

Figure S4. Longitudinal in-vivo imaging of nascent lung tumors. Relates to main figure 5.

Table S1. **Summary of Metacore GeneGO pathway analysis. Relates to main figure 4.**

## Acknowledgments

We express our gratitude to Peter Adams, Owen Sansom, David Bryant, Martin Eilers, Jennifer O’Neil & Ronan O’Hagan for helpful discussions over the course of this work; to Christine Kramer, Stephen Bell, Derek Miller and all the staff of the CRUK Beatson biological services unit for animal husbandry and assistance with animal protocols; to Peter Adams & Nikolay Pchelintsev for help establishing RNA-SEQ; and to all members of the Murphy lab for assistance in the preparation of this manuscript. Funding was provided by a MINT collaborative grant from Merck Sharpe & Dohme; Deutsche Krebshilfe grant 109220; EC FP7 Marie Curie Actions CIG 618448 “SERPLUC”; British Lung Foundation grants APHD13-5 and CSOBLF RG16-2 (all to DJM). B.K. was supported by EC H2020 Marie Curie actions mobility fellowship 705190 “NuSiCC”.

## References and Notes

1. Allemani, C., Weir, H. K., Carreira, H., Harewood, R., Spika, D., Wang, X. S., Bannon, F., Ahn, J. V., Johnson, C. J., Bonaventure, A., Marcos-Gragera, R., Stiller, C., Azevedo e Silva, G., Chen, W. Q., Ogunbiyi, O. J., Rachet, B., Soeberg, M. J., You, H., Matsuda, T., Bielska-Lasota, M., Storm, H., Tucker, T. C., Coleman, M. P. & Group, C. W. (2015) Global surveillance of cancer survival 1995-2009: analysis of individual data for 25,676,887 patients from 279 population-based registries in 67 countries (CONCORD-2), Lancet. 385, 977–1010.

2. Cancer Genome Atlas Research, N. (2014) Comprehensive molecular profiling of lung adenocarcinoma, Nature. 511, 543–50.

3. Ostrem, J. M., Peters, U., Sos, M. L., Wells, J. A. & Shokat, K. M. (2013) K-Ras(G12C) inhibitors allosterically control GTP affinity and effector interactions, Nature. 503, 548–51.

4. Lito, P., Solomon, M., Li, L. S., Hansen, R. & Rosen, N. (2016) Allele-specific inhibitors inactivate mutant KRAS G12C by a trapping mechanism, Science. 351, 604–8.

5. Roskoski, R., Jr. (2014) The ErbB/HER family of protein-tyrosine kinases and cancer, Pharmacological research. 79, 34–74.

6. Press, M. F., Cordon-Cardo, C. & Slamon, D. J. (1990) Expression of the HER-2/neu proto-oncogene in normal human adult and fetal tissues, Oncogene. 5, 953–62.

7. Prigent, S. A., Lemoine, N. R., Hughes, C. M., Plowman, G. D., Selden, C. & Gullick, W. J. (1992) Expression of the c-erbB-3 protein in normal human adult and fetal tissues, Oncogene. 7, 1273–8.

8. Bhattacharjee, A., Richards, W. G., Staunton, J., Li, C., Monti, S., Vasa, P., Ladd, C., Beheshti, J., Bueno, R., Gillette, M., Loda, M., Weber, G., Mark, E. J., Lander, E. S., Wong, W., Johnson, B. E., Golub, T. R., Sugarbaker, D. J. & Meyerson, M. (2001) Classification of human lung carcinomas by mRNA expression profiling reveals distinct adenocarcinoma subclasses, Proceedings of the National Academy of Sciences of the United States of America. 98, 13790–5.

9. Garber, M. E., Troyanskaya, O. G., Schluens, K., Petersen, S., Thaesler, Z., Pacyna-Gengelbach, M., van de Rijn, M., Rosen, G. D., Perou, C. M., Whyte, R. I., Altman, R. B., Brown, P. O., Botstein, D. & Petersen, I. (2001) Diversity of gene expression in adenocarcinoma of the lung, Proceedings of the National Academy of Sciences of the United States of America. 98, 13784–9.

10. Gollamudi, M., Nethery, D., Liu, J. & Kern, J. A. (2004) Autocrine activation of ErbB2/ErbB3 receptor complex by NRG-1 in non-small cell lung cancer cell lines, Lung cancer. 43, 135–43.

11. Chen, H. Y., Liu, C. H., Chang, Y. H., Yu, S. L., Ho, B. C., Hsu, C. P., Yang, T. Y., Chen, K. C., Hsu, K. H., Tseng, J. S., Hsia, J. Y., Chuang, C. Y., Chang, C. S., Li, Y. C., Li, K. C., Chang, G. C. & Yang, P. C. (2016) EGFR-activating mutations, DNA copy number abundance of ErbB family, and prognosis in lung adenocarcinoma, Oncotarget.

12. Zhang, J., Iwanaga, K., Choi, K. C., Wislez, M., Raso, M. G., Wei, W., Wistuba, II & Kurie, J. M. (2008) Intratumoral epiregulin is a marker of advanced disease in non-small cell lung cancer patients and confers invasive properties on EGFR-mutant cells, Cancer prevention research. 1, 201–7.

13. Sunaga, N., Kaira, K., Imai, H., Shimizu, K., Nakano, T., Shames, D. S., Girard, L., Soh, J., Sato, M., Iwasaki, Y., Ishizuka, T., Gazdar, A. F., Minna, J. D. & Mori, M. (2013) Oncogenic KRAS-induced epiregulin overexpression contributes to aggressive phenotype and is a promising therapeutic target in non-small-cell lung cancer, Oncogene. 32, 4034–42.

14. Citri, A. & Yarden, Y. (2006) EGF-ERBB signalling: towards the systems level, Nat Rev Mol Cell Biol. 7, 505–16.

15. Zhu, C. Q., da Cunha Santos, G., Ding, K., Sakurada, A., Cutz, J. C., Liu, N., Zhang, T., Marrano, P., Whitehead, M., Squire, J. A., Kamel-Reid, S., Seymour, L., Shepherd, F. A., Tsao, M. S. & National Cancer Institute of Canada Clinical Trials Group Study, B. R. (2008) Role of KRAS and EGFR as biomarkers of response to erlotinib in National Cancer Institute of Canada Clinical Trials Group Study BR.21, J Clin Oncol. 26, 4268–75.

16. de Bruin, E. C., Cowell, C., Warne, P. H., Jiang, M., Saunders, R. E., Melnick, M. A., Gettinger, S., Walther, Z., Wurtz, A., Heynen, G. J., Heideman, D. A., Gomez-Roman, J., Garcia-Castano, A., Gong, Y., Ladanyi, M., Varmus, H., Bernards, R., Smit, E. F., Politi, K. & Downward, J. (2014) Reduced NF1 expression confers resistance to EGFR inhibition in lung cancer, Cancer discovery. 4, 606–19.

17. Molina-Arcas, M., Hancock, D. C., Sheridan, C., Kumar, M. S. & Downward, J. (2013) Coordinate direct input of both KRAS and IGF1 receptor to activation of PI3 kinase in KRAS-mutant lung cancer, Cancer discovery. 3, 548–63.

18. Navas, C., Hernandez-Porras, I., Schuhmacher, A. J., Sibilia, M., Guerra, C. & Barbacid, M. (2012) EGF receptor signaling is essential for k-ras oncogene-driven pancreatic ductal adenocarcinoma, Cancer cell. 22, 318–30.

19. Ardito, C. M., Gruner, B. M., Takeuchi, K. K., Lubeseder-Martellato, C., Teichmann, N., Mazur, P. K., Delgiorno, K. E., Carpenter, E. S., Halbrook, C. J., Hall, J. C., Pal, D., Briel, T., Herner, A., Trajkovic-Arsic, M., Sipos, B., Liou, G. Y., Storz, P., Murray, N. R., Threadgill, D. W., Sibilia, M., Washington, M. K., Wilson, C. L., Schmid, R. M., Raines, E. W., Crawford, H. C. & Siveke, J. T. (2012) EGF receptor is required for KRAS-induced pancreatic tumorigenesis, Cancer cell. 22, 304–17.

20. Sun, C., Hobor, S., Bertotti, A., Zecchin, D., Huang, S., Galimi, F., Cottino, F., Prahallad, A., Grernrum, W., Tzani, A., Schlicker, A., Wessels, L. F., Smit, E. F., Thunnissen, E., Halonen, P., Lieftink, C., Beijersbergen, R. L., Di Nicolantonio, F., Bardelli, A., Trusolino, L. & Bernards, R. (2014) Intrinsic resistance to MEK inhibition in KRAS mutant lung and colon cancer through transcriptional induction of ERBB3, Cell reports. 7, 86–93.

21. Muthalagu, N., Neidler, S., Gyuraszova, K., Hedley, A., Hock, A., Braun, A., Nieswandt, Vousden, K., Dick, C., Sansom, O., Morton, J. & D.J. Murphy Dramatic acceleration of KRAS driven tumors by modest deregulation of MYC. Manuscript in preparation.

22. Subramaniam, D., He, A. R., Hwang, J., Deeken, J., Pishvaian, M., Hartley, M. L. & Marshall, J. L. (2015) Irreversible multitargeted ErbB family inhibitors for therapy of lung and breast cancer, Current cancer drug targets. 14, 775–93.

23. Junttila, M. R., Karnezis, A. N., Garcia, D., Madriles, F., Kortlever, R. M., Rostker, F., Brown Swigart, L., Pham, D. M., Seo, Y., Evan, G. I. & Martins, C. P. (2010) Selective activation of p53-mediated tumour suppression in high-grade tumours, Nature. 468, 567–71.

24. Feldser, D. M., Kostova, K. K., Winslow, M. M., Taylor, S. E., Cashman, C., Whittaker, C. A., Sanchez-Rivera, F. J., Resnick, R., Bronson, R., Hemann, M. T. & Jacks, T. (2010) Stage-specific sensitivity to p53 restoration during lung cancer progression, Nature. 468, 572–5.

25. Kerr, E. M., Gaude, E., Turrell, F. K., Frezza, C. & Martins, C. P. (2016) Mutant Kras copy number defines metabolic reprogramming and therapeutic susceptibilities, Nature. 531, 110–3.

26. Sarkisian, C. J., Keister, B. A., Stairs, D. B., Boxer, R. B., Moody, S. E. & Chodosh, L. A. (2007) Dose-dependent oncogene-induced senescence in vivo and its evasion during mammary tumorigenesis, Nature cell biology. 9, 493–505.

27. Moon, Y. W., Rao, G., Kim, J. J., Shim, H. S., Park, K. S., An, S. S., Kim, B., Steeg, P. S., Sarfaraz, S., Changwoo Lee, L., Voeller, D., Choi, E. Y., Luo, J., Palmieri, D., Chung, H. C., Kim, J. H., Wang, Y. & Giaccone, G. (2015) LAMC2 enhances the metastatic potential of lung adenocarcinoma, Cell Death Differ. 22, 1341–52.

28. Zhou, B. B., Fridman, J. S., Liu, X., Friedman, S. M., Newton, R. C. & Scherle, P. A. (2005) ADAM proteases, ErbB pathways and cancer, Expert Opin Investig Drugs. 14, 591–606.

29. Powell, C. A., Nasser, M. W., Zhao, H., Wochna, J. C., Zhang, X., Shapiro, C., Shilo, K. & Ganju, R. K. (2015) Fatty acid binding protein 5 promotes metastatic potential of triple negative breast cancer cells through enhancing epidermal growth factor receptor stability, Oncotarget. 6, 6373–85.

30. Ju, J. H., Oh, S., Lee, K. M., Yang, W., Nam, K. S., Moon, H. G., Noh, D. Y., Kim, C. G., Park, G., Park, J. B., Lee, T., Arteaga, C. L. & Shin, I. (2015) Cytokeratin19 induced by HER2/ERK binds and stabilizes HER2 on cell membranes, Cell Death Differ. 22, 665–76.

31. Gao, J., Aksoy, B. A., Dogrusoz, U., Dresdner, G., Gross, B., Sumer, S. O., Sun, Y., Jacobsen, A., Sinha, R., Larsson, E., Cerami, E., Sander, C. & Schultz, N. (2013) Integrative analysis of complex cancer genomics and clinical profiles using the cBioPortal, Sci Signal. 6, pl1.

32. Sanchez-Martin, M. & Pandiella, A. (2012) Differential action of small molecule HER kinase inhibitors on receptor heterodimerization: therapeutic implications, International journal of cancer Journal international du cancer. 131, 244–52.

33. Hock, A. K., Cheung, E. C., Humpton, T. J., Monteverde, T., Paulus-Hock, V., Lee, P., McGhee, E., Scopelliti, A., Murphy, D. J., Strathdee, D., Blyth, K. & Vousden, K. H. (2017) Development of an inducible mouse model of iRFP713 to track recombinase activity and tumour development in vivo, Sci Rep. 7, 1837.

34. Moll, H., Pranz, K., Musteanu, M., Grabner, B., Barbacid, M., Dome, B., Popper, H & E. Casanova Afatinib restrains KRAS driven lung tumorigenesis in

35. Cicchini, M., Buza, E. L., Sagal, K. M., Gudiel, A. A., Durham, A. C. & Feldser, D. M. (2017) Context-Dependent Effects of Amplified MAPK Signaling during Lung Adenocarcinoma Initiation and Progression, Cell reports. 18, 1958–1969.

36. Murphy, D. J., Junttila, M. R., Pouyet, L., Karnezis, A., Shchors, K., Bui, D. A., Brown-Swigart, L., Johnson, L. & Evan, G. I. (2008) Distinct thresholds govern Myc’s biological output in vivo, Cancer cell. 14, 447–57.

37. Grabocka, E., Pylayeva-Gupta, Y., Jones, M. J., Lubkov, V., Yemanaberhan, E., Taylor, L., Jeng, H. H. & Bar-Sagi, D. (2014) Wild-type H- and N-Ras promote mutant K-Ras-driven tumorigenesis by modulating the DNA damage response, Cancer cell. 25, 243–56.

38. Tebbutt, N., Pedersen, M. W. & Johns, T. G. (2013) Targeting the ERBB family in cancer: couples therapy, Nature reviews Cancer. 13, 663–73.

39. Jackson, E. L., Willis, N., Mercer, K., Bronson, R. T., Crowley, D., Montoya, R., Jacks, T. & Tuveson, D. A. (2001) Analysis of lung tumor initiation and progression using conditional expression of oncogenic K-ras, Genes & development. 15, 3243–8.

40. Church, D. M., Schneider, V. A., Graves, T., Auger, K., Cunningham, F., Bouk, N., Chen, H. C., Agarwala, R., McLaren, W. M., Ritchie, G. R., Albracht, D., Kremitzki, M., Rock, S., Kotkiewicz, H., Kremitzki, C., Wollam, A., Trani, L., Fulton, L., Fulton, R., Matthews, L., Whitehead, S., Chow, W., Torrance, J., Dunn, M., Harden, G., Threadgold, G., Wood, J., Collins, J., Heath, P., Griffiths, G., Pelan, S., Grafham, D., Eichler, E. E., Weinstock, G., Mardis, E. R., Wilson, R. K., Howe, K., Flicek, P. & Hubbard, T. (2011) Modernizing reference genome assemblies, PLoS Biol. 9, e1001091.

41. Kim, D., Pertea, G., Trapnell, C., Pimentel, H., Kelley, R. & Salzberg, S. L. (2013) TopHat2: accurate alignment of transcriptomes in the presence of insertions, deletions and gene fusions, Genome Biol. 14, R36.

42. Langmead, B. & Salzberg, S. L. (2012) Fast gapped-read alignment with Bowtie 2, Nat Methods. 9, 357–9.

43. Robinson, M. D., McCarthy, D. J. & Smyth, G. K. (2010) edgeR: a Bioconductor package for differential expression analysis of digital gene expression data, Bioinformatics. 26, 139–40.

44. Love, M. I., Huber, W. & Anders, S. (2014) Moderated estimation of fold change and dispersion for RNA-seq data with DESeq2, Genome Biol. 15, 550.

